# Incorporating metabolic activity, taxonomy and community structure to improve microbiome-based predictive models for host phenotype prediction

**DOI:** 10.1101/2023.01.20.524948

**Authors:** Mahsa Monshizadeh, Yuzhen Ye

## Abstract

We developed MicroKPNN, a prior-knowledge guided interpretable neural network for microbiome-based human host phenotype prediction. The prior-knowledge used in MicroKPNN includes the metabolic activities of different bacterial species, phylogenetic relationships, and bacterial community structure. Application of MicroKPNN to seven gut microbiome datasets (involving five different human diseases including inflammatory bowel disease, type 2 diabetes, liver cirrhosis, colorectal cancer, and obesity) shows that incorporation of the prior knowledge helped improve the microbiome-based host phenotype prediction. MicroKPNN outperformed fully-connected neural network based approaches in all seven cases, with the most improvement of accuracy in the prediction of type 2 diabetes. MicroKPNN outperformed a recently developed deep-learning based approach DeepMicro, which selects the best combination of autoencoder and machine learning approach to make predictions, in six out of the seven cases. More importantly, we showed that MicroKPNN provides a way for interpretation of the predictive models. Our results suggested that the metabolic potential of the bacterial species contributed more than the two other sources of prior knowledge. MicroKPNN is publicly available at https://github.com/mgtools/MicroKPNN.

## Introduction

The human gut microbiome play key roles in human health and diseases. Perturbations of the gut microbiota structure are associated with a variety of human diseases including cancer. Using microbial markers (genes, species, or pathways) that are differential between healthy individuals and patients, predictive models with promising accuracy have been built for predicting host phenotypes based on microbiome data [1, 2]. Gut microbiome composition was recently shown to be predictive of patient response to statins (the most common type of prescription drug which can lower cholesterol levels and reduce the risks of stroke and heart attack) and showed that Bacteroides-enriched individuals have a higher risk of statin-induced metabolic disruption [3].

Microbiome-based human host phenotype prediction has benefited from the recent advances in Machine Learning (ML) and Artificial Intelligence (AI) algorithms. SIAMCAT is a machine learning toolbox developed to address the issues related to ML algorithms in microbiome studies such as poor generalization [4]. Goallec et al. [2] showed that the prediction accuracy depended on the choice of ML algorithms and types of metagenomic data, and presented a computational framework for inferring microbiome-derived features for host phenotype predictions. This paper [5] demonstrated the benefit of building multi-disease models to achieve accurate microbiome-based predictive models for human phenotype prediction.

Deep learning methods including various autoencoders were also exploited for learning the representation of quantitative microbiome profile in a lower dimensional latent space, which were then used for building predictive models for host disease prediction [6]. DeepMicro [6] took advantage of recently developed autoencoders, including shallow autoencoder (SAE), deep autoencoder (DAE), variational autoencoder (VAE), and convolutional autoencoder (CAE), to achieve a low dimensional representation from high-dimensional microbiome profile. Various machine learning classification algorithms (SVM, random forest, and MLP) were then applied on the learned representation for prediction. DeepMicro was shown to perform well as tested on six different disease datasets. However, the results showed that the performance of the various autoencoders varied.

Considering that microbial species are phylogenetically related, there are a few attempts that tried to incorporate the phylogenetic relationship in the deep learning models for microbiome-based prediction. Ph-CNN [7] and PopPhy-CNN [8] are two of such approaches that share similar core ideas to represent species or OTU abundance profiles in 2D matrices such that the phylogenetic relationship among the species/OTUs are retained to some extent in the 2D matrics. These two approaches differ in how they achieve this goal. Ph-CNN uses the patristic distance between species/OTUs (computed from the tree) together with a sparsified version of multidimensional scaling to embed the phylogenetic tree in a Euclidean space. PopPhy-CNN prepares the 2D matrix representing the phylogenetic tree populated with the relative abundance of microbial taxa in a metagenomic sample, such that, for a given row, the children of the nodes from that row are selected and their abundances are placed in the subsequent row in the order that their parents appear, starting with the left-most column. EPCNN [9] is a more recent effort that uses an ensemble strategy utilizing different microbial features and taxonomic representation for microbiome-based prediction, aiming to reduce overfitting.

Despite the success of applying deep learning approaches to build microbiome-based predictors, the downside of these algorithms is the lack of interpretability due to their black-box nature. To overcome this issue, we designed a neural network architecture (MicroKPNN) that incorporates various microbial relationships (metabolic, phylogenetic, and community) in the model to improve the performance of microbiome-based prediction and interpretability of the models. Using this prior-knowledge guided neural network, we can examine which microbial relationship plays important roles in the prediction of host health status.

Our MicroKPNN is inspired by the KPNN approach [10], a deep learning approach for interpretable deep learning and biological discovery and designed to predict human cell state from single cell RNA-seq data in deep neural networks that are constructed based on biological knowledge. In KPNN each node corresponds to a human protein or a gene, and each edge corresponds to a regulatory relationship that has been documented in biological databases. A notable difference between MicroKPNN and KPNN is that MicroKPNN uses a shallow neural network, with only one hidden layer, allowing a more straightforward interpretation of predictions and examination of the importance of prior knowledge and microbial relationship for prediction.

Gut microbes interact among themselves and with hosts. Various metabolic activities in the gut attribute to the interactions. The gut microbiota makes an important contribution to human metabolism by contributing enzymes that are not encoded by the human genome, for example, the breakdown of polysaccharides and polyphenols, and the synthesis of vitamins [11]. Gut microbes can also break down host-derived substrates such as mucins. Among microbial community members, competitive relationships may be formed if microbes compete for the same resources, and they may also form cooperation relationships via metabolic cross-feeding in a shared environment [12, 13]. Comparison of predicted metabolic-interactions and species co-occurrence patterns suggested that habitat-filtering shapes the gut microbiome [12]. Using literature mining, Sung et al [14] curated an interspecies network of the human gut microbiota (called NJS16) comprising hundreds of microbial species and three human cell types metabolically interacting through >4,400 small-molecule transport (import or export) and macromolecule degradation events. Metabolic activities have been used for providing explanations for example observed differential species/genes, however, they haven’t been explicitly used in predictive models.

Microbial organisms form communities. Metagenomic co-occurrence has been widely applied in metagenomic studies to construct microbiome networks and better understand microbiome community structures [15–17]. We have recently inferred networks of microbial networks using metagenomic co-occurrence approach, taking advantage of the availability of many gut metagenomic sequencing datasets derived from healthy and diseased individuals, and recent methodology advances in network inference that can deal with sparse compositional data [18]. From the networks, communities of microbes were identified. In this paper, we ask if such community information can be utilized to improve microbiome-based host phenotype prediction.

We developed MicroKPNN, to incorporate metabolic activities and community information of gut microbes, in addition to their phylogenetic relationship, in constrained neural networks for microbiome-based prediction of host phenotype. We tested MicroKPNN on seven microbiome datasets derived from cohorts of individuals with different phenotypes and compared its performance with other approaches. We showed that it achieved comparable performance across all datasets with the state of the art phenotype predictors including deep learning approaches DeepMicro and EPCNN which involve much more complex network architectures. Remarkably, MicroKPNN achieved even better accuracy than the existing approaches for diseases including liver cirrhosis, IBD, and obesity. We showed that due to the shallow nature of our models, and the fact that hidden nodes carry biological meanings, MicroKPNN provides interpretability that was missed by the existing deep learning approaches.

## Methods

### Collection of the human gut metagenomic samples

We first used the same six datasets as those used for developing ML models including MetAML [19], DeepMicro [6] and EPCNN [9]. Table 1 summarizes the six datasets and the diseases they represent. For fair comparison, we used the species profile abundance data downloaded from the DeepMicro [6] GitHub repository at https://github.com/minoh0201/DeepMicro. The species abundance profiles for these six datasets were estimated by MetaPhlAn2 [20]. The species-level relative abundance profile consists of real values in [0,100] representing the percentages of the species in the total observed species for a sample; the numbers sum up to 100 (%) for each sample.

**Table 1:**
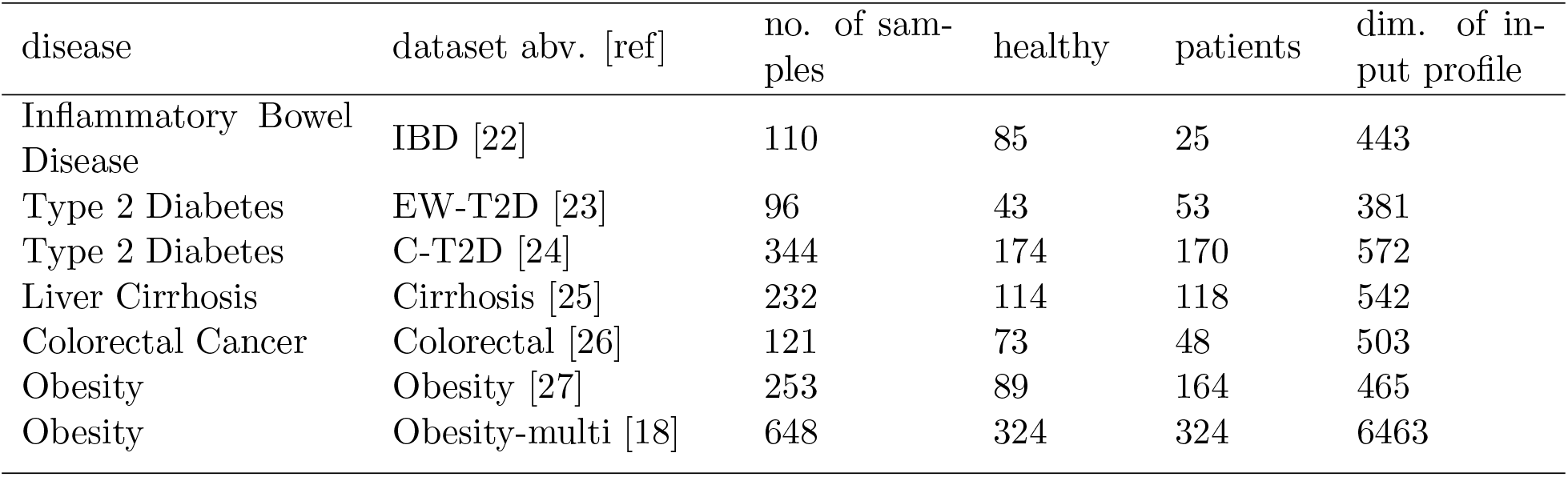
Summary of human gut microbiome datasets used for disease state prediction.

In addition, we used metagenomic datasets associated with obesity from 15 studies in an attempt to showcase the application of our tool (this collection is referred to as Obesity-multi in Table 1). The metagenome datasets were downloaded from NCBI SRA and were analyzed using the Kraken+Bracken approach [21] to derive taxonomic assignments and quantification. See more details of the datasets and data processing in [18]. The list of datasets and their abundance profiles are available in the MicroKPNN github repository (see Implementation below).

### Network structure

The main idea of MicroKPNN is to use prior knowledge to constrain the links between the nodes in the neural network, similar to the knowledge-primed neural networks (KPNN) method [10]. Different from KPNN, MicroKPNN is a neural network that is constructed according to the prior knowledge of the bacterial species to improve host phenotype prediction using microbiome data. In MicroKPNN, hidden nodes have biological meanings, such as taxa, and every edge has a relation interpretation. This network offers insights into not only the importance of individual microbial species and the consequent impact of the microbiota on the host, but also the importance of taxonomic and metabolic variations relevant to host phenotypes.

Specifically, MicroKPNN uses a neural network that has three layers (see Figure 1): the input layer, one hidden layer, and the output layer. Our results (see Results) show that such a shallow network doesn’t sacrifice good performance compared to the deep learning approaches that have been developed for microbiome-based prediction. The input layer is species abundance derived from human gut microbiome samples. The hidden layer includes four different groups of nodes (shown in different colors in Figure 1):

- Metabolites. This part of the network architecture encompasses the relationships among gut microbial species and chemical compounds. Since microbial species have different metabolic capabilities with some being the producers of certain metabolites and others being the consumers of certain metabolites, each metabolite may be represented as two nodes in the hidden layer. One node for the metabolite has edges coming from the producer species in the input layer, whereas the other node has edges coming from the consumer species. For example, there are two nodes of L-lactate in the hidden layer (L-lactate consumption and L-lactate production), and there are edges connecting the producers (such as *Bacteroides ovatus*) with L-lactate production, and there are edges connecting the consumers (such as *Acetobacter pasteurianus*) with L-lactate consumption. In addition, many bacterial species can degrade macromolecules including mucin (mucus glycoprotein) and cellulose, and edges will be added to connect the bacterial species and the corresponding macromolecules in the hidden layer. For example, many microbes including *Bacteroides fragilis* can degrade mucin and utilize it as a nutrient source for growth [28, 29], and therefore edges will be created connecting the mucin-degraders and mucin.
- Taxa. MicroKPNN uses a much simpler approach for encoding the phylogenetic relationship as compared to the previous deep-learning approaches that use phylogenetic information (such as Ph-CNN). The NCBI hierarchical taxonomy [30] is used to encode the taxonomic relationship. For example, if *Bacteroides* (a genus) is a node in the hidden layer, all the species in the input layer that belong to this genus will have an edge to this hidden node. MicroKPNN will try different taxonomic ranks including genus, order, class, and phylum (a hyperparameter in the model), and the rank that results in the best performance will be selected.
- Communities. Each node represents a community, and all species that are part of the community have an edge connecting to the node. The communities were computationally inferred using the Leiden algorithm [31] from a species co-occurrence network [18].
- Pure hidden nodes. Unlike the nodes in the above three groups that have corresponding biological meanings, these unknown nodes are added to alleviate the potential problem of losing information due to incomplete prior knowledge. This part of the network is fully connected, i,e., there is a link between every input node and every hidden node in this group. The number of unknown hidden nodes is another hyperparameter in our model.

**Figure 1:**
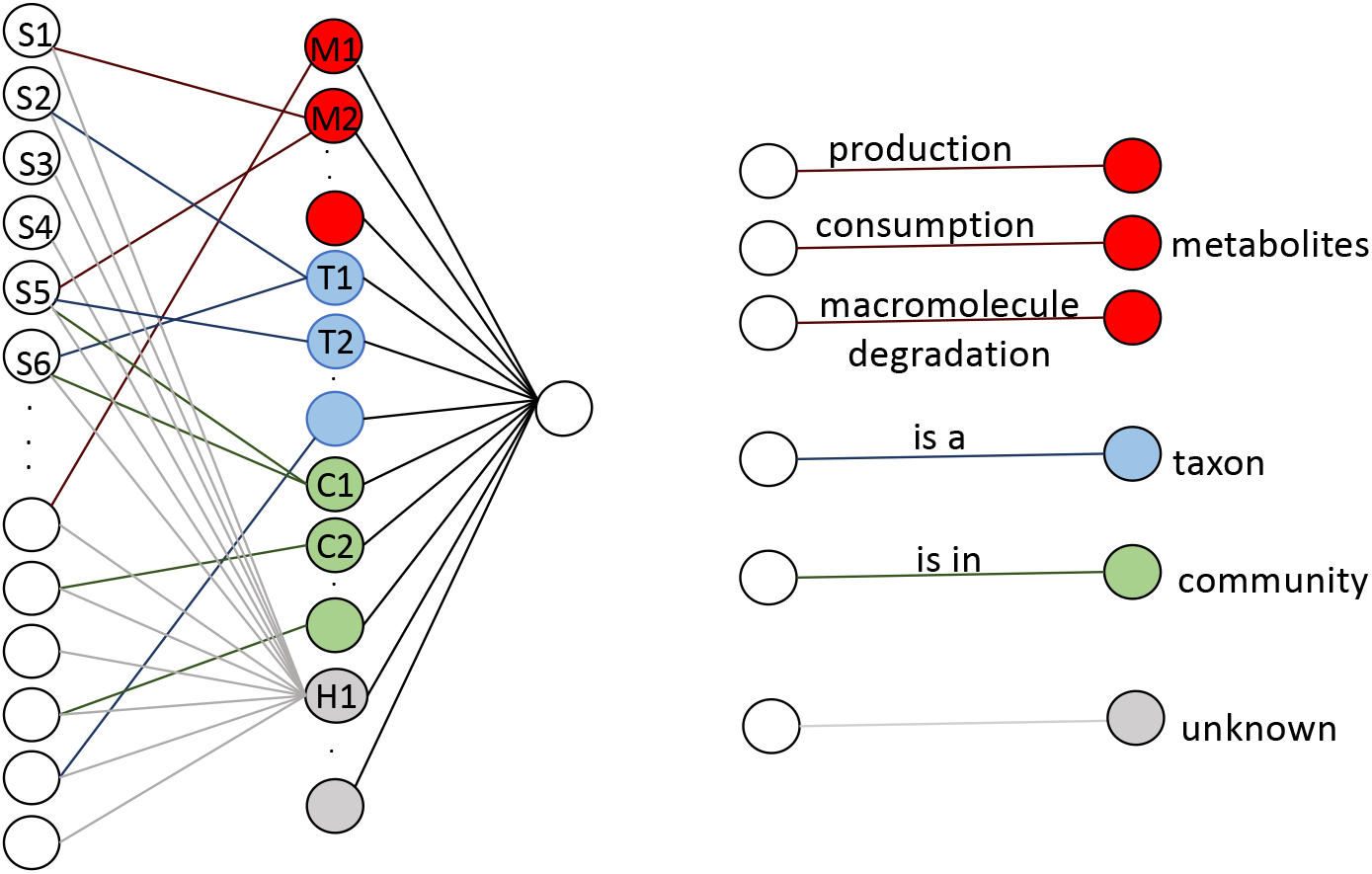
The neural network structure used in MicroKPNN. It is composed of three layers (shown on the left). In the input layer, each node is a species, and the hidden layer includes nodes of four different groups: metabolites (red), taxa (blue), communities (green), and unknown hidden nodes (gray). The links between the input nodes and the nodes in the hidden layer represent different biological meanings (shown on the right).

### Implementation

We adapted KPNN for the training and application of the neural network for microbiome-based host phenotype prediction. KPNN workflow was implemented in python (3.7.13), using TensorFlow for NN training. Here are the settings for training in KPNN: edge weights randomly initialized, a sigmoid activation function for all hidden and output nodes, and a weighted cross-entropy with L2 regularization as the loss function. We added Python scripts that can be used to prepare the network constraints to KPNN for microbiome applications. The network constraints are encoded using a list of edges between the species in the input layer and the nodes in the hidden layer: metabolic edges are created according to the NJS16 metabolic network [14], community edges are inferred based on the network file (in the standard gml format) of microbial communities [18], and the taxonomic edges are created according to the NCBI taxonomy. For clarity, we called our adopted version of KPNN for microbiome-based prediction as MicroKPNN, which is available as a GitHub repository at https://github.com/mgtools/MicroKPNN.

#### Fully-connected neural networks without using prior knowledge

To show the importance of including the prior knowledge as the constraints for prediction in MicroKPNN, we also implemented fully connected neural networks (fc-NNs) without using the prior knowledge for comparison. We used two implementations of fully connected NNs: our implementation of NN using the Keras library [32], and MicroKPNN including only the unknown nodes in the hidden layer. We note the difference between the two implementations is that the latter has extra processes including early stopping and dropout that are inherited from KPNN. For fair comparison, in fully connected neural networks, the number of hidden nodes is set to be the as same as the total number of hidden nodes (including the hidden nodes with biological meanings and the pure hidden nodes) in the best performing setting of MicroKPNN for each dataset, resulting in a consistent number of hidden nodes across MicroKPNN and fully-connected NNs for each prediction.

### Training, evaluation, and interpretation

For each dataset, we split data into training, validation, and test sets in the ratio of 6:2:2 with a given random partition seed, keeping the ratio between classes in both training and test set to be the same as that of the given dataset. There are two hyperparameters in MicroKPNN, the taxonomic rank and the number of unknown hidden nodes (the number of metabolite nodes and the number of community nodes are fixed). We tested the following taxonomic ranks: kingdom, phylum, class, order, family, or genus. For unknown hidden nodes, we tried 10, 20, 30, 40, 50, 60, 70, 80, 90 and 100 for each dataset. We ran the MicroKPNN algorithm five times on each of the different combinations of hidden nodes to compute the average accuracy and standard error. The area under the receiver operating characteristics curve (AUC) was used for performance evaluation.

The predictive models once trained were analyzed to calculate node weights as a reflection of node importance for the predictions. KPNN applied small perturbations to each hidden node separately and measured changes in network output, thus quantifying the importance of each node to the output of the network. Because the sign of the resulting node weights is largely arbitrary, KPNN uses the absolute value of the node weights as a measure of the importance of each node in the trained KPNNs. Similarly in MicroKPNN, we used the absolute value of the node weights to quantify the importance of the nodes in the hidden layer. Since we used 5 repeated calculations, we used the average of the importance scores from the repeated runs to quantify the importance of the nodes.

## Results

### Optimization of the neural network structure for the different diseases

We used different combinations of taxonomic ranks and the number of hidden nodes (the hyperparameters) to see which setting resulted in the most accurate predictions. Table 2 summarizes the best performance of the MicroKPNN on the seven datasets and the corresponding configuration of the NN architecture. For example, the best NN predictor trained on EW-T2D contains 309 total nodes in the hidden layer, including 34 nodes representing different orders of the bacterial species, and 10 unknown hidden nodes. Figure 2 shows the impacts of the different parameters on the performance of the prediction of T2D (EW-T2D). MicroKPNN gave an almost perfect prediction for cirrhosis as shown in Table 2, suggesting the significant differences in the bacterial composition of the cirrhosis patients compared to healthy controls.

**Figure 2:**
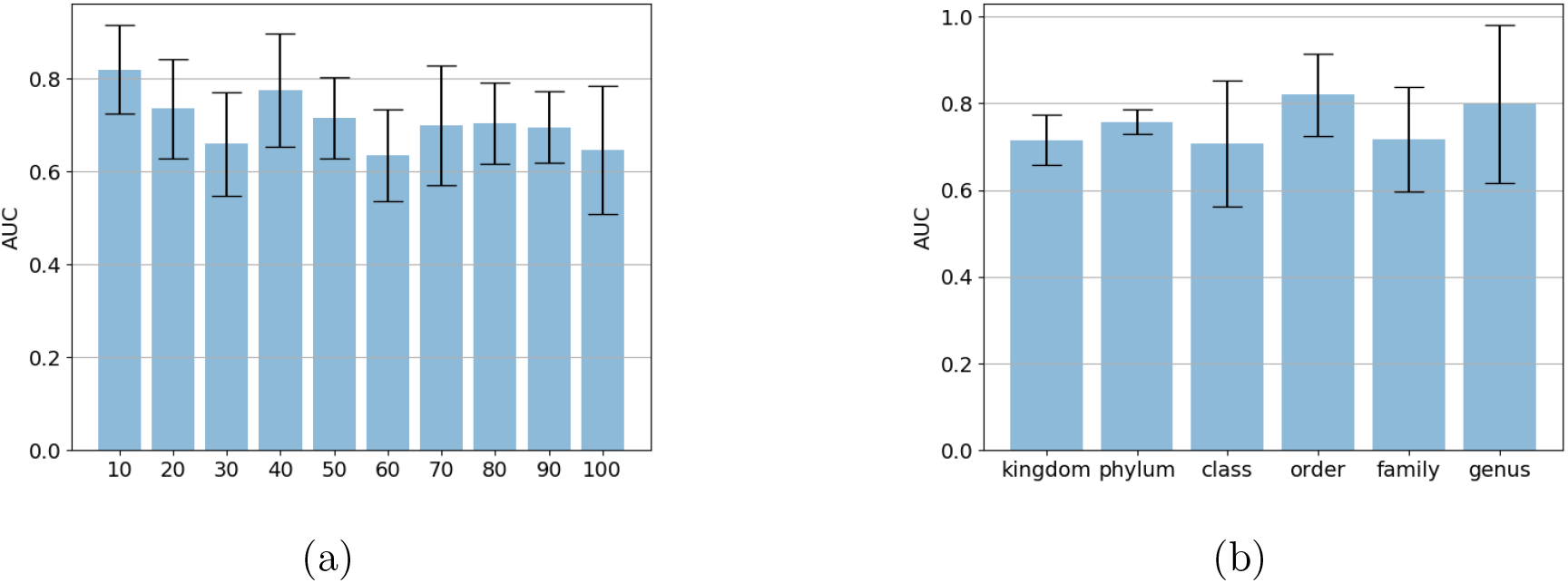
Impacts of the hyperparameters on the MicroKPNN performance for the EW-T2D dataset. (a) the number of unknown hidden nodes (taxonomic rank set to “order”); (b) taxonomic rank (the number of unknown hidden nodes set to 10).

Table 2 also shows that different taxonomic ranks have different impacts on the performance of the predictors, depending on the diseases. For example, using taxonomic information at the order level gave the best performance for T2D prediction (EW-T2D), whereas for obesity, using genus in the hidden layer gave the best prediction comparing to other taxonomic ranks. Although there is no single taxonomic rank that performed the best across all different diseases, genus in general, gave a relatively good performance, with average AUCs of 0.953 (IBD), 0.797 (EW-T2D), 0.734 (C-T2D), 0.955 (Cirrhosis), 0.837 (Colorectal), 0.699 (Obesity), and 0.829 (Obesity-multi).

**Table 2:** Summary of best performing neural network architecture for each dataset and their average AUC.

| dataset | no. of nodes in different groups in the hidden layer |  |  |  |  | avg. AUC |
| --- | --- | --- | --- | --- | --- | --- |
|  | all | taxon (rank) | metabolite | community | unknown |  |
| IBD | 519 | 176 (genus) | 234 | 29 | 80 | 0.953 |
| EW-T2D | 309 | 34 (order) | 234 | 31 | 10 | 0.820 |
| C-T2D | 365 | 27 (class) | 240 | 38 | 60 | 0.753 |
| Cirrhosis | 354 | 40 (order) | 239 | 35 | 40 | 0.969 |
| Colorectal | 403 | 65 (family) | 234 | 34 | 70 | 0.846 |
| Obesity | 479 | 184 (genus) | 233 | 32 | 30 | 0.699 |
| Obesity-multi | 822 | 190 (order) | 276 | 336 | 20 | 0.875 |

### Comparison of MicroKPNN with fully-connected NN and existing deep learning predictors

Table 3 shows the comparison of MicroKPNN with the fully-connected NN (without the guide of prior knowledge) and two existing state-of-the-art deep learning approaches for microbiome-based prediction (DeepMicro and EPCNN). MicroKPNN achieved drastic performance improvements comparing to NN without constraining the network connection according to the prior knowledges (i.e., fully-connected NN with the same number of nodes in the hidden layer as the corresponding MicroKPNN) on four out of the seven datasets: IBD, EW-T2D, Colorectal, and Obesity (the improvement on the T2D, Cirrhosis, and Obesity-multi dataset were modest).

**Table 3:** Comparison of MicroKPNN with different methods including NNs that are fully connected (fc-NN) in averaged AUC and standard deviation (in parenthesis).

| Dataset | MicroKPNN | fc-NN (keras) | fc-NN (KPNN) | DeepMicro<sup>a</sup> | EPCNN<sup>c</sup> |
| --- | --- | --- | --- | --- | --- |
| IBD | **0.953** (0.019) | 0.865 (0.031) | 0.678 (0.089) | 0.873 (0.067) | NA<sup>d</sup> |
| EW-T2D | 0.820 (0.047) | 0.595 (0.065) | 0.580 (0.062) | **0.829** (0.087) | 0.789 (0.056) |
| C-T2D | 0.753 (0.013) | 0.675 (0.033) | 0.723 (0.019) | 0.725 (0.056) | **0.813** (0.024) |
| Cirrhosis | **0.969** (0.009) | 0.823 (0.022) | 0.947 (0.021) | 0.888 (0.025) | 0.953 (0.007) |
| Colorectal | 0.846 (0.020) | 0.624 (0.038) | 0.764 (0.053) | 0.809 (0.103) | **0.906** (0.013) |
| Obesity | **0.699** (0.045) | 0.539 (0.023) | 0.608 (0.022) | 0.674 (0.076) | NA |
| Obesity-multi | **0.875** (0.017) | 0.820 (0.014) | 0.826 (0.045) | 0.763 (0.042)<sup>b</sup> | NA |

MicroKPNN outperformed DeepMicro and EPCNN, two of the most recent deep learning ML approaches, on four out of the seven datasets (IBD, Cirrhosis, Obesity, Obesity-multi), and achieved worse, but still reasonable AUCs on the other three datasets (EW-T2D, C-T2D, and Colorectal). MicroKPNN outperformed DeepMicro in six out of the seven cases (except EW-T2D). We note DeepMicro tries different representation deep learning approaches and ML algorithms (RF, SVM and MLP) and reports the best AUCs. For the Obesity-multi dataset, we ran DeepMicro using all combinations of autoencoders and ML algorithms using the same input to MicroKPNN as the input, and the combination of SAE and RF resulted in the best performance (results shown in Table 2). We also note that DeepMicro may either use species abundance or gene abundance as the input, and here we focus on comparison with DeepMicro using species abundance. EPCNN were based on predictors also trained using species abundances information, but there are differences: EPCNN uses both known and unknown species and the quantification was achieved using Micropro [33]. Both DeepMicro and EPCNN were based on complex deep learning models: DeepMicro used the various autoencoders (SAE, DAE, VAE, and CAE), and EPCNN used multiple convolution layers. By contrast, MicroKPNN’s model is a much simpler neural network with only one hidden layer, and in combination with the use of nodes with different biological meanings in the hidden layer (which provide good interpretability of the models see below), we believe MicroKPNN’s performance is very encouraging.

MicroKPNN’s accuracy for the Obesity dataset is the lowest among all datasets, however, its AUC (0.699) is significantly greater than the AUCs achieved by other approaches, including DeepMicro using species abundance profile as the input (AUC=0.674) (DeepMicro achieved an AUC of 0.650 when it used the gene profile instead of species profile as the input for prediction). The results suggest that the microbiome difference is more subtle between healthy controls and patients with obesity comparing to other phenotypes.

The predictor trained using datasets from multiple studies (Obesity-multi) achieved much more accurate predictions of obesity comparing to the predictor built from a single study (Obesity). We attribute the improvement (AUC of 0.875 vs 0.699) to using more datasets from multiple studies, among others (e.g., using a different approach for taxonomic assignment and quantification). It also suggests that when datasets from different studies become available, it is beneficial to include all of them to improve the accuracy of the predictive models and perhaps make the predictive models more generalizable. For comparison, DeepMicro’s average AUC is 0.763 on this dataset.

### Intepretability of MicroKPNN

We used the node weights as measures of the importance of the corresponding nodes (metabolic activity, taxonomic rank, community of species, and unknown hidden nodes) and therefore to provide an explanation of the impacts of the inputs (the species) on the prediction. By examining the importance scores of the hidden nodes, we could compare the contribution of the different groups of nodes to the prediction. Figure 3 shows the boxplots of the importance scores of the most important nodes in each group for the different diseases. The comparison shows that the metabolite nodes contributed more than the taxon nodes and the community nodes across all diseases. The notable example is cirrhosis prediction, in which the metabolite nodes contribute obviously more than the community and taxon nodes to the performance (see Figure 3e).

**Figure 3:**
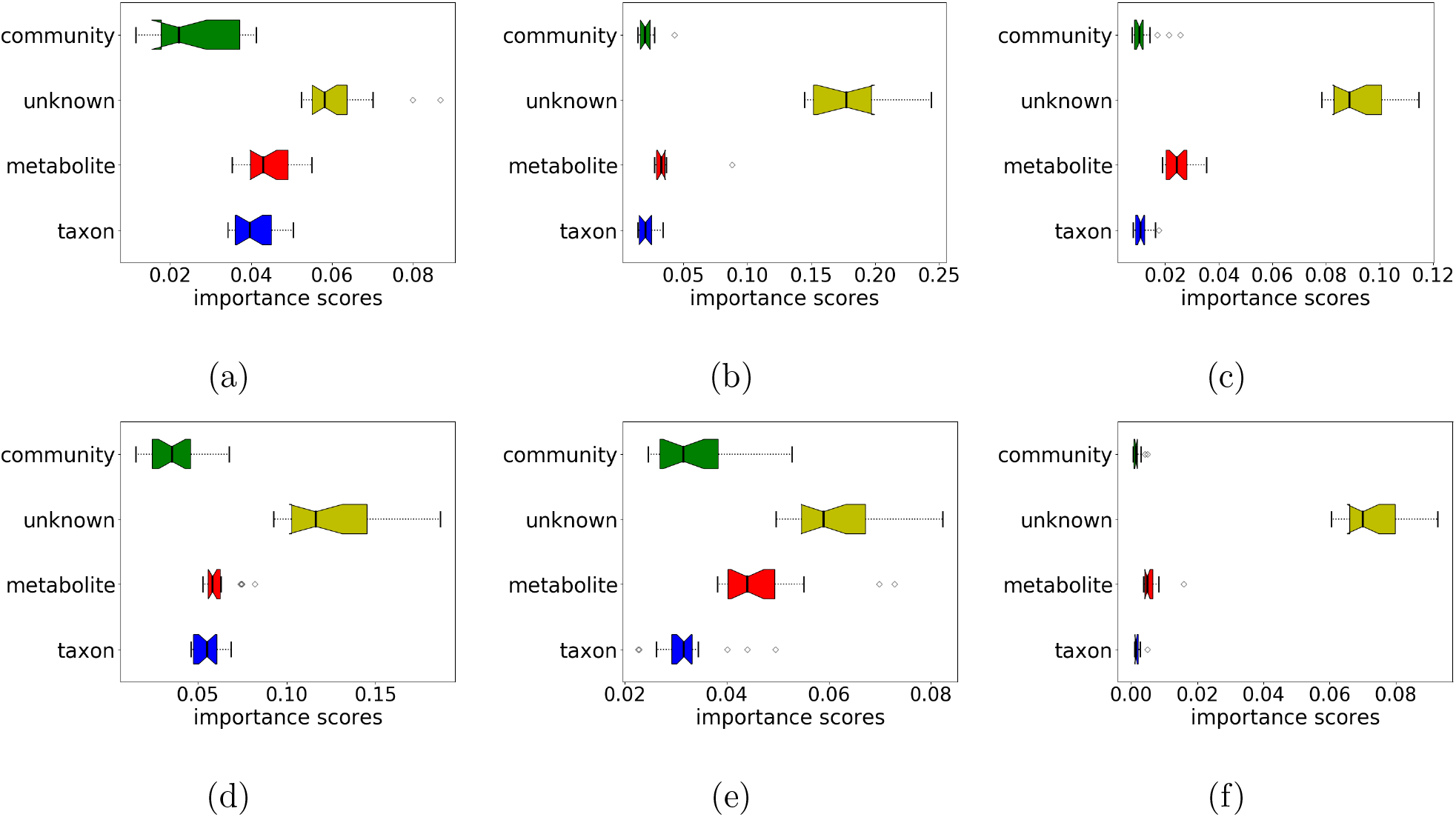
Contributions of the different groups of hidden nodes to the prediction as measured by importance scores. (a) IBD; (b) EW-T2D; (c) C-T2D; (d) Obesity; (e) Cirrhosis; (f) Colorectal Cancer.

The importance scores of the individual nodes in the hidden layer also provide a way for suggesting biologically meaningful explanations to the microbiome-based predictors. Figure 4a shows the top five nodes for each group in the hidden layer that are important for the prediction of liver cirrhosis based on microbiome data. Our prediction model showed that L-Idonate, CO2, and mucin glycoprotein were the top three most important metabolite nodes that contributed to the prediction. Among the bacterial species that are involved in mucin glycoprotein degradation (i.e., mucin consumers), we observed that *Ruminococcus gnavus* was highly elevated in cirrhosis patients. Increase of *R. gnavus* was found to be implicated in the degradation of elements from the mucus layer providing an explanation for the impaired intestinal barrier function and systematic inflammation in LC patients [34]. Lactate consumption and production were also picked up as important nodes by MicroKPNN, suggesting the importance of the bacterial species that produce and/or digest these metabolites. It is well known that bacteria produce intermediate fermentation products including lactate, but these are normally detected at low levels in feces from healthy individuals due to extensive utilization of them by other bacteria [11, 35]. Among the taxon nodes, *Bifidobacteria* had the highest importance score; previous studies have shown that patients with chronic liver disease have varying degrees of intestinal microflora imbalance with a decrease of total Bifidobacterial counts [36].

**Figure 4:**
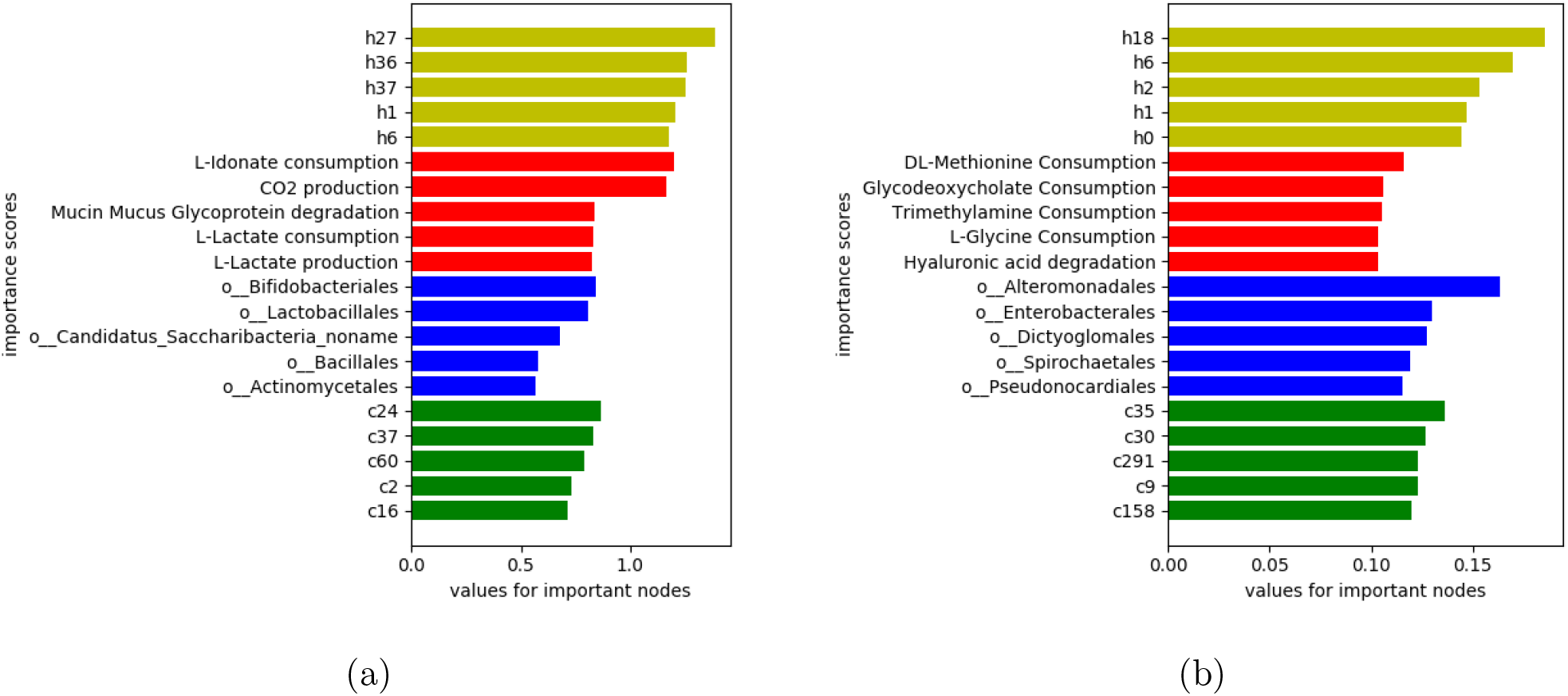
a) Importance scores of the hidden nodes for microbiome-based host phenotype predictions. The top five most important nodes of each group in the hidden layer for prediction of cirrhosis and obesity (trained on Obesity-multi with datasets from multiple studies) are shown in (a) and (b), respectively. The bars are highlighted in different colors: yellow for unknown nodes, red for metabolites, blue for taxa, and green for communities.

Figure 4b shows the contribution of different hidden nodes to the prediction of obesity (trained using Obesity-multi dataset). Among the metabolite nodes, Methionine consumption was picked by MicroKPNN as the most important node. It was shown that dietary methionine restriction increased fat oxidation in obese adults with metabolic syndrome [37], and our results suggested the importance of studying bacterial consumers of methionine consumption when studying obesity. The bacterial community that contributed the most to the prediction is c35 which is composed of four Actinobacteria (*Actinomadura* sp. NAK00032, *Nonomuraea* sp. ATCC 55076, *Streptomyces* sp. WAC 01438, *Egicoccus halophilus*) and two Proteobacteria (*Pseudoxanthomonas spadix* and *Geobacter bremensis*).

## Discussion

MicroKPNN uses a simple architecture, but by leveraging on prior knowledge of microbial species, it provides promising predictions of host phenotype based on microbiome composition as shown on all seven datasets. Comparison of the importance scores of different prior knowledge showed that the metabolic activities had the largest impact on the performance of predictions. The difference between the relative importance scores of the hidden nodes with that of the unknown nodes indicates the knowledge gap between the microbial species and their interaction with human hosts. For colorectal cancer, it was mostly the unknown nodes that contributed to the prediction, indicating that although the predictor has a good AUC of 0.846, the existing knowledge about the metabolic potential and bacterial interactions has limited value for interpreting the prediction of colorectal cancer based on the microbiome.

The predictive models we built in this work are based on species abundance. It has been shown (including our own work) that using bacterial genes typically (not always) results in better predictive models [2, 38]. Nevertheless, our knowledge-primed MicroKPNN’s was able to fill in some of the gaps between species-based and gene-based predictions. For example, among the seven datasets that we tested, DeepMicro using gene markers as the inputs significantly outperformed DeepMicro based on species abundance for IBD prediction (AUC=0.955 vs AUC=0.873) and Cirrhosis (AUC=0.940 vs AUC=0.888) [6]. MicroKPNN achieved comparable performance as DeepMicro using gene markers for IBD prediction, with AUC=0.953 and even outperformed DeepMicro using gene markers for cirrhosis prediction (AUC=0.969) (see the comparison in Table 3). A future direction of our work is to expand MicroKPNN so that it can take gene abundance as the input for microbiome-based prediction. We anticipate that more complex architecture will need to be adopted to incorporate the prior-knowledge, for example, multiple layers of hidden nodes to capture the hierarchical relationship of the genes.

